# Miniaturization technologies for cost-effective AmpliSeq library preparation for next generation sequencing

**DOI:** 10.1101/467464

**Authors:** Eri Ogiso-Tanaka, Akito Kaga, Makita Hajika

## Abstract

**Purpose:** AmpliSeq technology, the target enrichment method for next-generation sequencing (NGS), enables quick and easy detection of the genomic “hot spot” region frequently mutated in species. Even though the cost of NGS has decreased, library preparation cost accounts for a more significant proportion of the total cost. If AmpliSeq library can be prepared at a lower cost, large-scale precision oncology can be more easily carried out. Furthermore, this technology can be widely applied not only to medical research, but also to polymorphism detection in biology. This study aimed to reduce the cost of AmpliSeq library preparation by adopting miniaturization technology.

**Methods:** We used approximately 10 ng of genomic DNA for ultra-multiplex PCR of 384, 768, 1152, 1920, and 3072 amplicons. Multiplex PCR was performed in a total volume of 1.6, 2.0, and 2.4 μL, using a nano-liter liquid handler, for library preparation.

**Results:** The success rate of library construction decreased with decreasing total multiplex PCR reaction volume. Using 1.6-, 2.0-, and 2.4-μL reactions, the success rates of ultra-multiplex PCR were 25%, 95%, and 100%, respectively. We could stably create libraries of the correct amplicon size, with an amplicon number of approximately 1500 or less. As a result of NGS, uniformity of PCR amplification and read length of quality-checked libraries were hardly affected by the number of amplicons.

**Conclusion:** Here, we show that the minimum volume for a stable reaction was 2.4 μL and the maximum number of amplicons obtained was approximately 1500. The protocol saved 86.8% in reagent usage and reduced handling time by 85% compared to that required by the manual protocol. Therefore, miniaturization technologies could reduce the cost of AmpliSeq library preparation through minimization of reagents.

## Introduction

Genotyping by sequencing (GBS), using next-generation sequencing (NGS), is becoming an increasingly useful technique in the fields of biology and agronomy [1-3]. In recent years, the cost of sequencing per base has drastically reduced, although the library preparation cost and time involved still remain high.

AmpliSeq technology (Thermo Fisher Scientific, Waltham, MA) is a GBS method that is often used in cancer research [4]. Although the AmpliSeq library kit is very expensive (about 64 USD or 9,500 JPY/sample), highly multiplexed PCR (of approximately 6000 primer pairs) is made possible with it. AmpliSeq technology is beginning to be used in other fields as well and is sold as AgriSeq in the fields of biology and agronomy [5, 6].

In the field of agronomy, especially in crop-breeding sites, genotyping of thousands of large samples is carried out often. However, GBS is still expensive, requiring a considerable amount of time and labor for analysis. It is difficult to apply it for practical crop breeding, and thus, is mainly used in research or project. A quick way to decrease the cost of GSB is to decrease library preparation cost. The simplest way for that is to reduce the reaction volume. In manual library preparation, volume reduction by more than half to one quarter is not realistic. Quality deterioration of the library is highly probable with reduction in the amount of reaction volume in case of manual preparation. Therefore, to sufficiently reduce the cost, it is necessary to have a robot capable of transferring minute quantities of liquid that cannot otherwise be handled by humans.

Till date, only few protocols for miniaturized library preparation in GBS have been reported. To reduce the cost and time limitations of current library preparation techniques, we tried to adapt our AmpliSeq, one such GBS method that follows a miniaturized reagent protocol, using Mosquito HTS (TTP Labtech, Royston, UK), which is a positive displacement pipetting instrument. Mosquito HTS offers highly accurate and precise multichannel pipetting (8 or 16 channels) from 25 nL to 1.2 μL. Mosquito HTS was originally used for protein crystallization [7]. Recently, Mosquito HTS and another model Mosquito HV have been used for single-cell RNASeq analysis [8]. Mosquito HV being a high-volume model (500 nL to 5 μL) does not save enough reagent and does not minimize the cost of AmpliSeq library preparation. Here, we describe a method that allows miniaturized, automated, and cost- and time-efficient 384-well library preparation with its quality and performance.

## Materials

### DNA preparation

DNA was extracted from young leaves of four soybean cultivars (cv. Aso Masari, cv. Suzuotome, cv. Tachiyutaka, and cv. Toyoshirome) using bead-based method (chemagic DNA plant kits, PerkinElmer, Waltham, MA) with BioSprint 96 DNA plant kit on robotic workstation (Qiagen, Hilden, Germany). Extracted DNA quality was evaluated based on DIN value (DNA Integrity Number), which is an index showing the fragmentation extent of DNA using TapeStation 4200 (Agilent Technologies, Santa Clara, CA). DNA concentration was measured with the Qubit Fluorometer (Thermo Fisher Scientific, Waltham, MA) by exciting the sample at 485 nm and measuring the fluorescence intensity at 520 nm. The instrument was calibrated with the Quant-iT Qubit dsDNA BR and HS Assay kit (Thermo Fisher Scientific, Waltham, MA), following the manufacturer’s instructions.

### Amplicon primers

In this study, 3072 AmpliSeq primer pairs were used [6]. Among them, 96, 192, 288, 384, 768, 1152, 1536, 1920, 2304, 2688, and 3072 primer sets (panel) were mixed with 100 nM of each primer. These primer sets were chosen sequentially from 3072 amplicon panels with the smallest amplicon number. The in-silico amplicon size distributions are shown in Figure 1. The peak size in all panels was approximately 280 bp (Figure 1).

**Figure 1.**
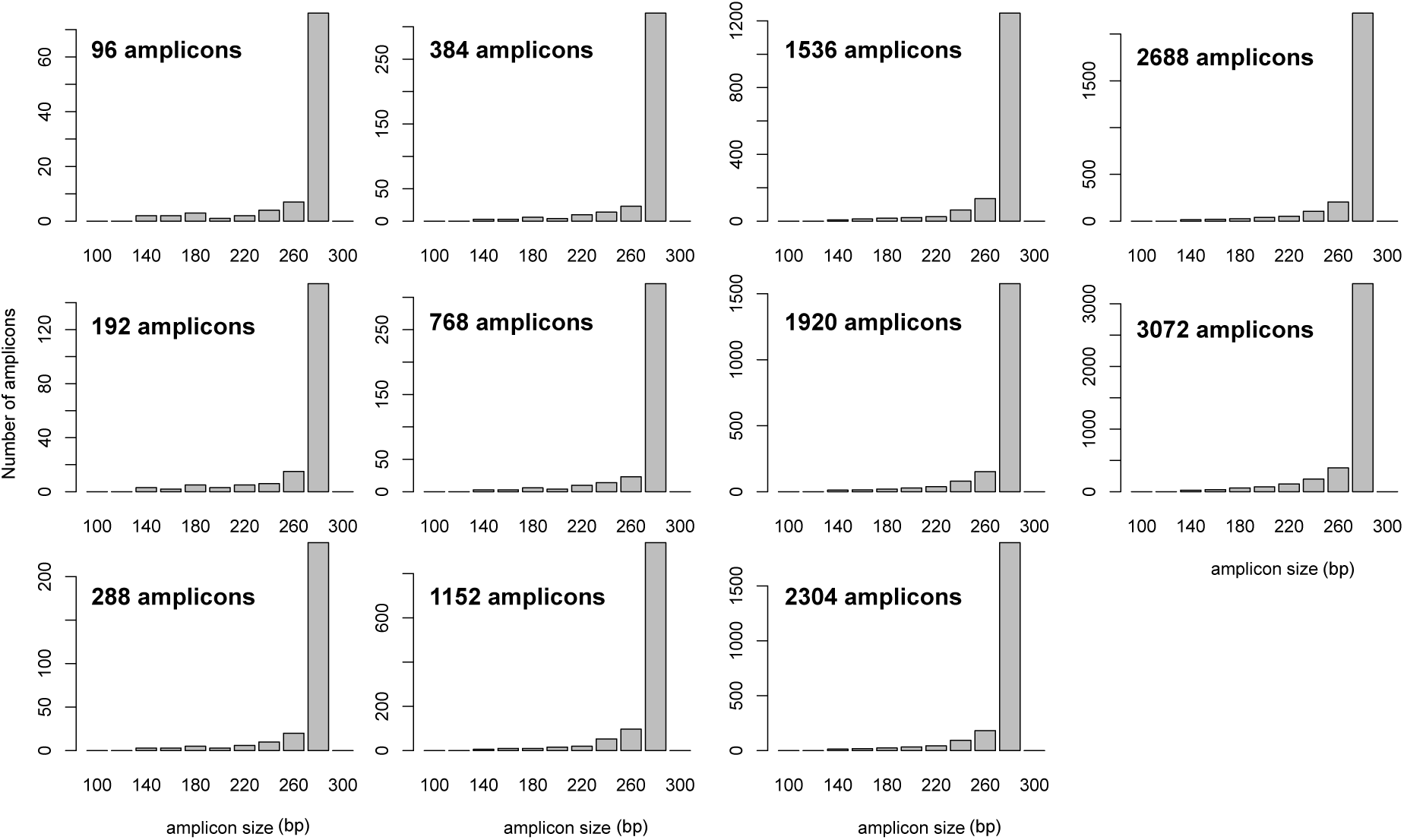
Distribution of amplicon size on each panel

## Results and Methods

### AmpliSeq library preparation with Mosquito HTS

NGS library was constructed using the Ion AmpliSeq Library Kit 2.0 (Thermo Fisher Scientific, Waltham, MA) [4]. To miniaturize this procedure for Mosquito HTS, we tested a modified protocol in reduced volume of reagent (Table 1). Although making a master mix reduces the number of dispensing steps, a dead volume is likely to occur. Therefore, template DNA and each reagent were manually dispensed into each row of 384-well Low Volume Serial Dilution (LVSD) plates [source plate] for automated liquid handling by Mosquito HTS (TTP Labtech, Royston, UK), and each plate was stored at −20 ºC thereafter. Multiplex PCR was performed using Hard-Shell PCR 384-well Plates (Bio-rad, Richmond, CA) with MicroAmp Clear Adhesive Film (Applied Biosystems). When general PCR plates (MicroAmp Optical 384-well Reaction Plate, ABI: Applied Biosystems, Foster City, CA) were used, they were found to bend in heat during PCR causing the needle to hit the bottom to stop Mosquito HTS at the next dispense step for FuPa.

**Table 1.**
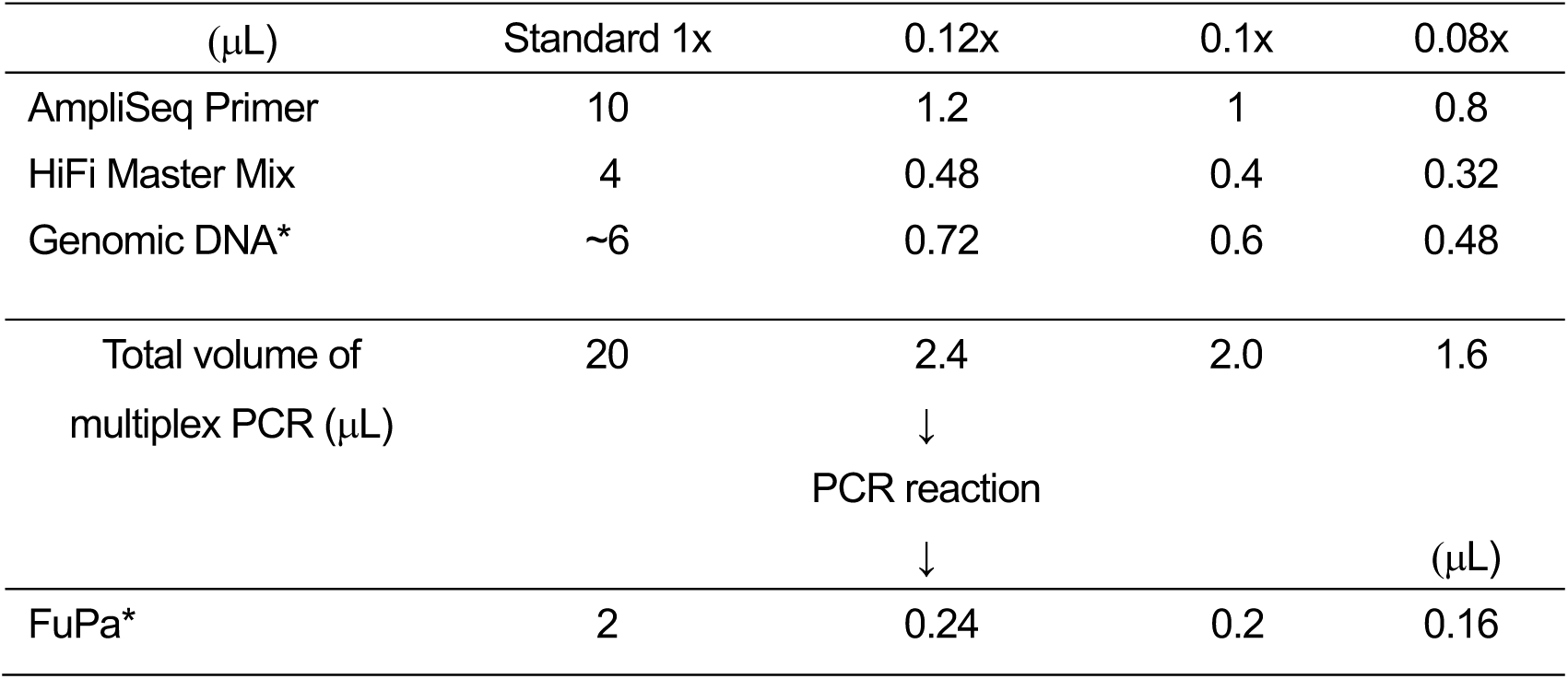
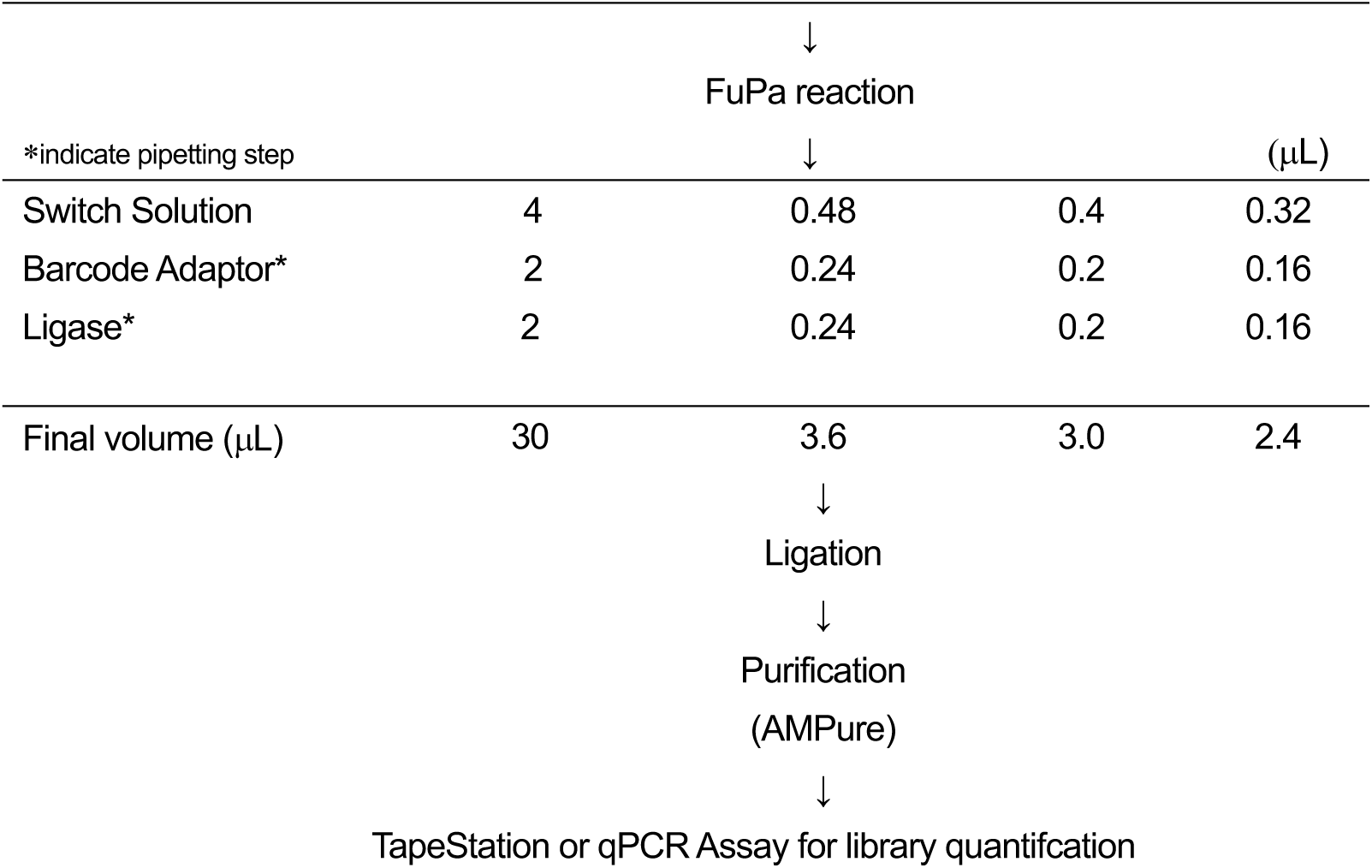
Standard and miniaturization protocols for AmpliSeq library preparation

For multiplex PCR amplification, 10 ng of each genomic DNA sample (cv. Williams 82) was amplified using 1 primer pool (96, 192, 288, 384, 768, 1152, 1536, 1920, 2304, 2688, and 3072 amplicons panel) per reaction (Table 1) [6]. The multiplex PCR was performed in 2.4 μL (12% volume) using individual GeneAmp PCR System 9700 (ABI). Since the maximum capacity of Mosquito HTS is 1.2 μL, the total reaction volume was set at 2.4 μL with the amount of 2x AmpliSeq primer set to 1.2 μL.

In four samples (cv. Aso Masari, cv. Suzuotome, cv. Tachiyutaka, cv. Toyoshirome), multiplex PCR was performed using 1 primer pool (384, 768, 1152, 1920, and 3072 amplicons panel) in 2.4 μL (12% volume), 2.0 μL (10%), and 1.6 μL (8%) using the best performing GeneAmp PCR System 9700 (ABI) and 3072 amplicons in 20 μL (100% volume) using SimpliAmp Thermal Cycler (ABI). In the reaction volume for miniaturized PCR, evaporation and vapor loss are critical. However, the MicroAmp Clear Adhesive Film (Applied Biosystems) could prevent vapor loss without Vapor-Lock or mineral oil. The reaction mix was heated for 2 min at 99 ºC for enzyme activation, followed by 16 (384–3071 amplicons) or 20 (96–288 amplicons) two-step cycles at 99 ºC for 15 s and 60 ºC for 4 min (96-1536 amplicons) or 8 min (1920-3072 amplicons), ending with a holding period at 10 ºC. The amplified samples were digested using 0.24, 0.2, or 0.16 μL FuPa enzymes per sample at 55 ºC for 10 min, followed by enzyme inactivation at 60 ºC for 20 min. To enable multiple libraries to be loaded per chip, 0.24, 0.2, or 0.16 μL of a unique diluted mix, including IonCode768 Barcode and Ion P1 Adapters (Thermo Fisher Scientific, Waltham, MA), was ligated to the end of the digested amplicons using 0.24, 0.2, or 0.16 μL of DNA ligase at 22 ºC for 1 h, followed by ligase inactivation for 10 min at 72 ºC. The library volume was made up to 10 μL with low TE.

Barcoded libraries from Williams 82 were confirmed by D1000 ScreenTape with Agilent 4200 TapeStation (Agilent Technologies, Santa Clara, CA) according to the manufacturer’s instruction. All 2.4-μL multiplex PCR reactions with 96–3072 amplicon panels were successful, but the library size did not stabilize when the number of amplicons exceeded 1920 (Figure 2).

**Figure 2.**
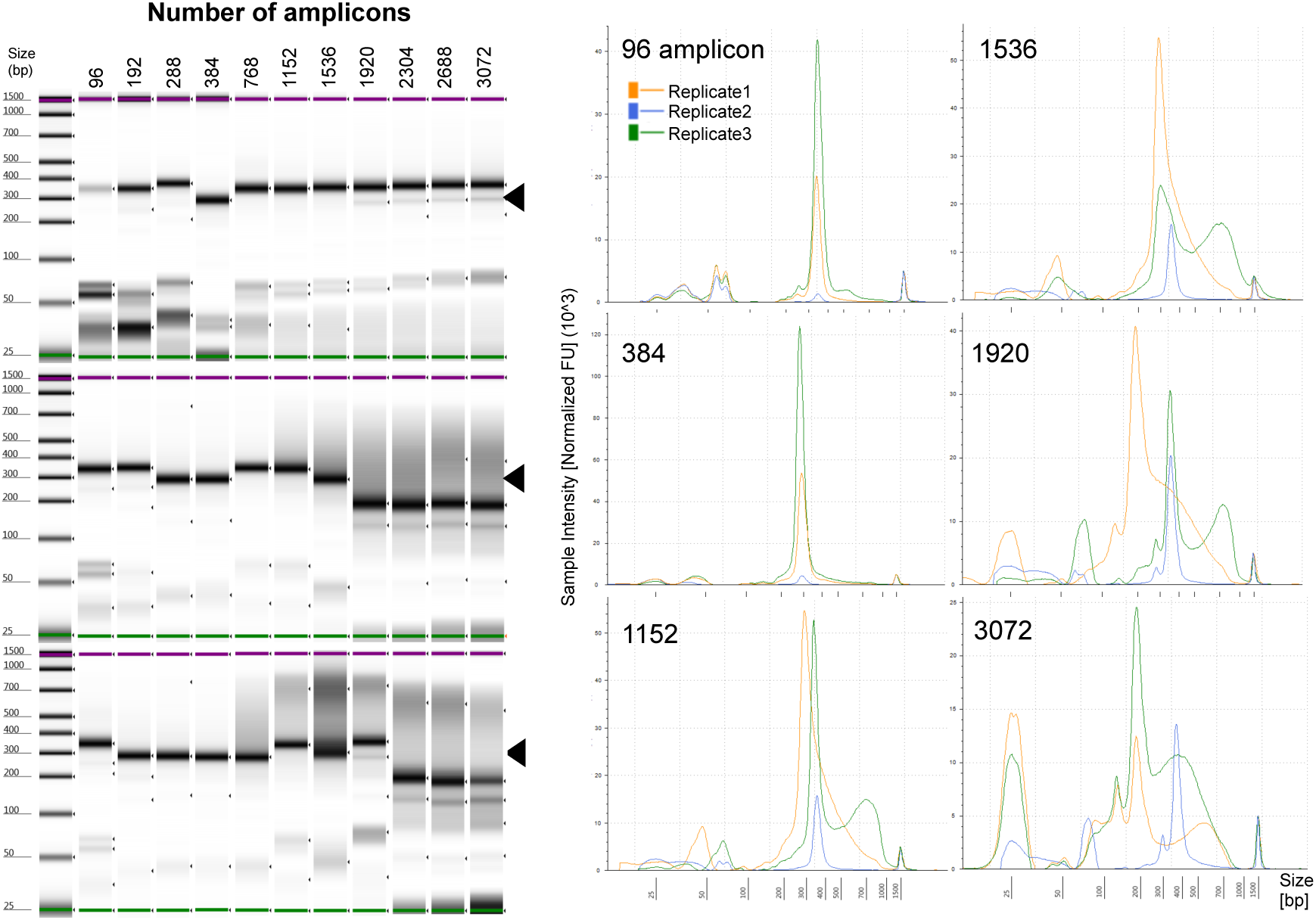
TapeStation 4200 profiles of AmpliSeq libraries after ligation reaction. (A) Image obtained by digital electrophoresis of individual replicates (replicate 1-3). Black arrow indicates 300-bp size. Lower-size band (< 100 bp) corresponds to primer or primer dimer. Sizing accuracy is ± 10%. Left side indicates electronic ladder. Sizing accuracy for analysis with electronic ladder is ± 20%. (B) The graphic shows the library size distribution for PCR products of 96, 384, 1152, 1536, 1920, and 3072 amplicon panels in replicates 1-3 (A). These multiplex PCRs were performed in different PCR thermal cyclers (GeneAmp PCR System 9700, ABI).

Although all dispensing steps utilized Mosquito HTS nano-liter handling in this protocol, the Mosquito could not be utilized for the magnetic bead-cleaning steps in this process. Therefore, we performed bead cleaning and size selection using Agencourt AMPure XP beads (Beckman Coulter, Brea, CA) by a manual approach. We used 1.2x (12 μL) of AMPure XP beads for bead-cleaning, followed by the addition of 40 μL of freshly prepared 70% ethanol to each miniaturized library. We usually used 1.5x of AMPure XP in the standard protocol (20 μL library volume) or half volume (10 μL) for AmpliSeq library prep [4], but the primer and a small-sized product (< 100 bp) remained in the miniaturized protocol. A SPRIPlate 384 Magnet Plate (Beckman Coulter) was used to minimize library elution volume. We repeated the washing step twice, completely removed the ethanol, and air-dried the beads for 3–5 min (may be judged by the alcoholic smell), while the plate was on the magnetic rack. The library was eluted from the beads with 12-μL low TE, and 10 μL of the supernatant was transferred to a clean plate.

To check the concentration of libraries prepared by the miniaturized protocol, barcoded libraries were quantified by qPCR using the Ion Library Quantitation Kit. This was performed using 2.4 μL of 2x TaqMan Master Mix, 0.24 μL of 20x Ion TaqMan Assay, and diluted libraries. The libraries were diluted 500- and 1000-fold (200- and 100-nL libraries in 100 μL low TE), and 2.16 μL of diluted libraries was transferred to 384-well PCR plate (MicroAmp Optical 384-Well Reaction Plate with Barcode, ABI) in preparation for TaqMan qPCR. These reagents were also dispensed by the Mosquito HTS (each library and TaqMan Master Mix was dispensed twice since the upper-limit volume of Mosquito HTS was 1.2 μL). Five dilutions of an *Escherichia coli* DH10B Ion Control Library of known concentration were run in the same plate in triplicate, as a standard. Following qPCR, the concentration of each library was calculated using their Ct values in a linear regression of Ct vs LOC, generated using the standards. Thereafter, each library was mixed at equal concentration according to the library concentration and number of amplicons.

The mixed adapter-ligated libraries were purified using 1.5-fold volume of AMPure XP Reagent, followed by the addition of 150 μL of freshly prepared 70% ethanol to each library. The washing step was repeated twice, the ethanol was completely removed, and beads were air-dried for 3–5 min, while the plate was on the magnetic rack. The library was eluted from the beads with 23 μL low TE; 20 μL of the supernatant was transferred to a clean tube.

The concentration and size of amplicons were determined using D1000 ScreenTape with Agilent 4200 TapeStation (Agilent Technologies, Santa Clara, CA) according to the manufacturer’s instruction. After quantification, mixed library was diluted to a concentration of 100 pM prior to template preparation.

Next, the barcoded libraries of four samples with 384, 768, 1152, 1920, and 3072 amplicon panels in 2.4 (12% volume), 2.0 (10%), and 1.6 (8%) μL reaction volumes were finally made up to 10 μL by adding low TE, and libraries purified by AMpure XP were confirmed by qPCR. Although the concentration of the libraries was 7–236 nM, 16 libraries (one in 2.0 μL reaction volume and 15 in 1.6 μL reaction volume) had a concentration of 0 nM (Table 2). The success rate of library construction was 100% (20/20) in the 2.4-μL reaction volume similar to the rate in the 20-μL volume, but was 95% (19/20) and 25% (5/20) in the 2.0-μL and 1.6-μL reaction volumes, respectively. The failure of library preparation occurred independently of the number of amplicons and DNA samples (Table 2), indicating that DNA quality or the number of amplicons did not account for the failure in library preparation. In samples in which library preparation failed, PCR products were not detected by the TapeStation 4200 system (data not shown), indicating that the multiplex PCR did not work well in miniaturized PCR reaction volumes of 2 μL or less.

**Table 2.** The concentration of library by qPCR-based library quantitation (nM).

| Sample name | Number of amplicons | 20 μL | 2.4 μL | 2.0 μL | 1.6 μL |
| --- | --- | --- | --- | --- | --- |
| Tachiyutaka | 382 | - | 68 | 7 | n.d. |
|  | 768 | - | 27 | 20 | n.d. |
|  | 1152 | - | 57 | 18 | n.d. |
|  | 1920 | - | 95 | 35 | n.d. |
|  | 3072 | 202 | 54 | 40 | n.d. |
| Asomasari | 382 | - | 17 | 8 | 7 |
|  | 768 | - | 13 | n.d. | n.d. |
|  | 1152 | - | 16 | 21 | n.d. |
|  | 1920 | - | 56 | 34 | n.d. |
|  | 3072 | 176 | 52 | 48 | n.d. |
| Toyoshirome | 382 | - | 8 | 6 | n.d. |
|  | 768 | - | 50 | 174 | n.d. |
|  | 1152 | - | 11 | 62 | n.d. |
|  | 1920 | - | 35 | 39 | 14 |
|  | 3072 | 236 | 49 | 39 | n.d. |
| Suzuotome | 382 | - | 15 | 12 | 8 |
|  | 768 | - | 53 | 148 | 15 |
|  | 1152 | - | 45 | 14 | n.d. |
|  | 1920 | - | 74 | 29 | 18 |
|  | 3072 | 132 | 43 | 28 | n.d. |
| The success rate of library construction |  | (4/4)<br>100% | (20/20)<br>100% | (19/20)<br>95% | (5/20)<br>25% |

n.d. : Not detected

The quality-checked libraries were pooled into the appropriate concentration, and the mixed library was diluted to a concentration of 100 pM prior to template preparation as described above.

### Sequencing and coverage analysis

Final libraries were sequenced on the IonTorrent S5 system (Thermo Fisher Scientific, Waltham, MA). Template preparation consisting of emulsion PCR, enrichment of beads containing the template, and chip loading, was performed with the Ion Chef instrument and Ion S5 Kit-Chef according to the manufacturer’s instruction (Thermo Fisher Scientific, Waltham, MA). After the preparation of ion sphere particles (ISPs), sequencing for 500 cycles was performed with an Ion Torrent Ion S5 system using Ion 520 and 540 Chip (Thermo Fisher Scientific, Waltham, MA) according to the manufacturer’s instruction. The sequence data was mapped to the soybean genome reference version 2.0 (Gmax275: http://genome.jgi.doe.gov/pages/dynamicOrganismDownload.jsf?organism=Phytozome#, downloaded on May 15, 2015) by Ion Torrent Suite v5.8.0 software. The software was optimized for Ion Torrent raw data analysis—alignment of Torrent Mapping Alignment Program (TMAP) v5.8.17 and coverage analysis v5.8.0.8 plugin.

We obtained an average depth coverage of 875 (a total of 18.8 M reads) in Williams 82 with 96, 192, 288, 384, 768, 1152, 1536, 1920, 2304, 2688, and 3072 amplicon panels. On target rate ranged from 56.13% (1152 amplicon panel) to 85.84% (384 amplicon panel), and uniformity of coverage ranged from 51.09% (1152 amplicon panel) to 93.59% (768 amplicon panel) (Figure 3). Although there was almost no difference in mean read length between amplicon panels, the on-target rate was low at 1152 and 1536 amplicon panels. Uniformity tended to be lower with amplicons over 1152 (Figure 3).

Among a total of 19,442,437 reads obtained, 18,792,850 (96.7%) were mapped to a reference genome using TMAP (Figure 3). On-target rate represents the percentage of reads mapped to any targeted region relative to all reads mapped to the reference. Uniformity represents the percentage of bases in all targeted regions covered by at least 0.2x the average base read depth.

**Figure 3.**
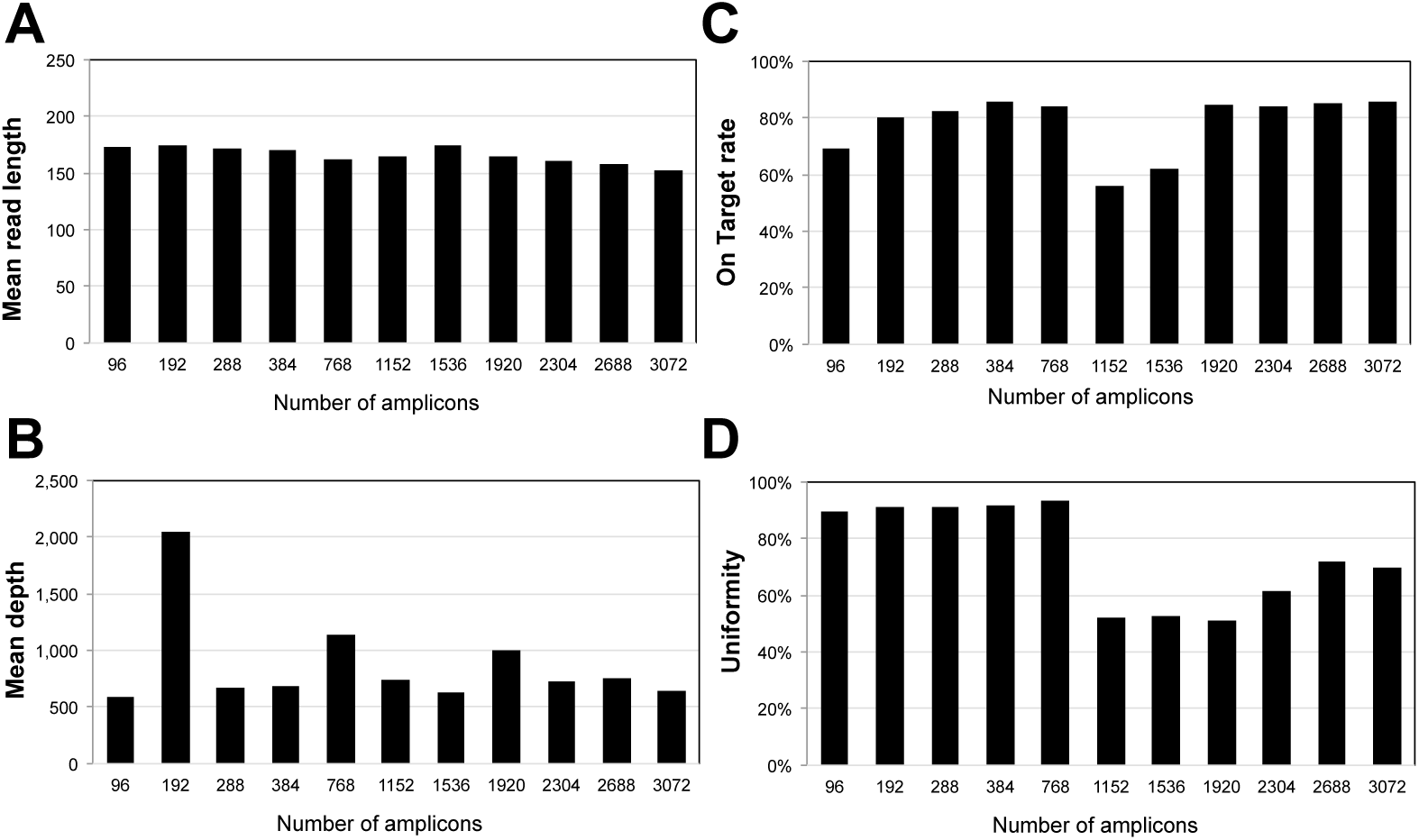
Sequencing result of Williams 82 in each amplicon panel. (A) On-target rate, (B) Mean depth, and (C) Uniformity.

We obtained an average depth coverage of 672 (a total of 65.9 M reads) in four samples with 384, 768, 1152, 1920, and 3072 amplicon panels. A similar trend was observed for a low % of on-target rate and uniformity in the 1152 amplicon panel. These results indicated that the differences in on-target rate and uniform PCR amplification are panel-dependent and that miniaturized volumes of reagents do not affect on-target rate or uniformity. Compared to that obtained with the standard 20-μL volume, mean read length, on-target rate, and uniformity of the miniaturized protocol were not different (Figure 3).

**Figure 3.**
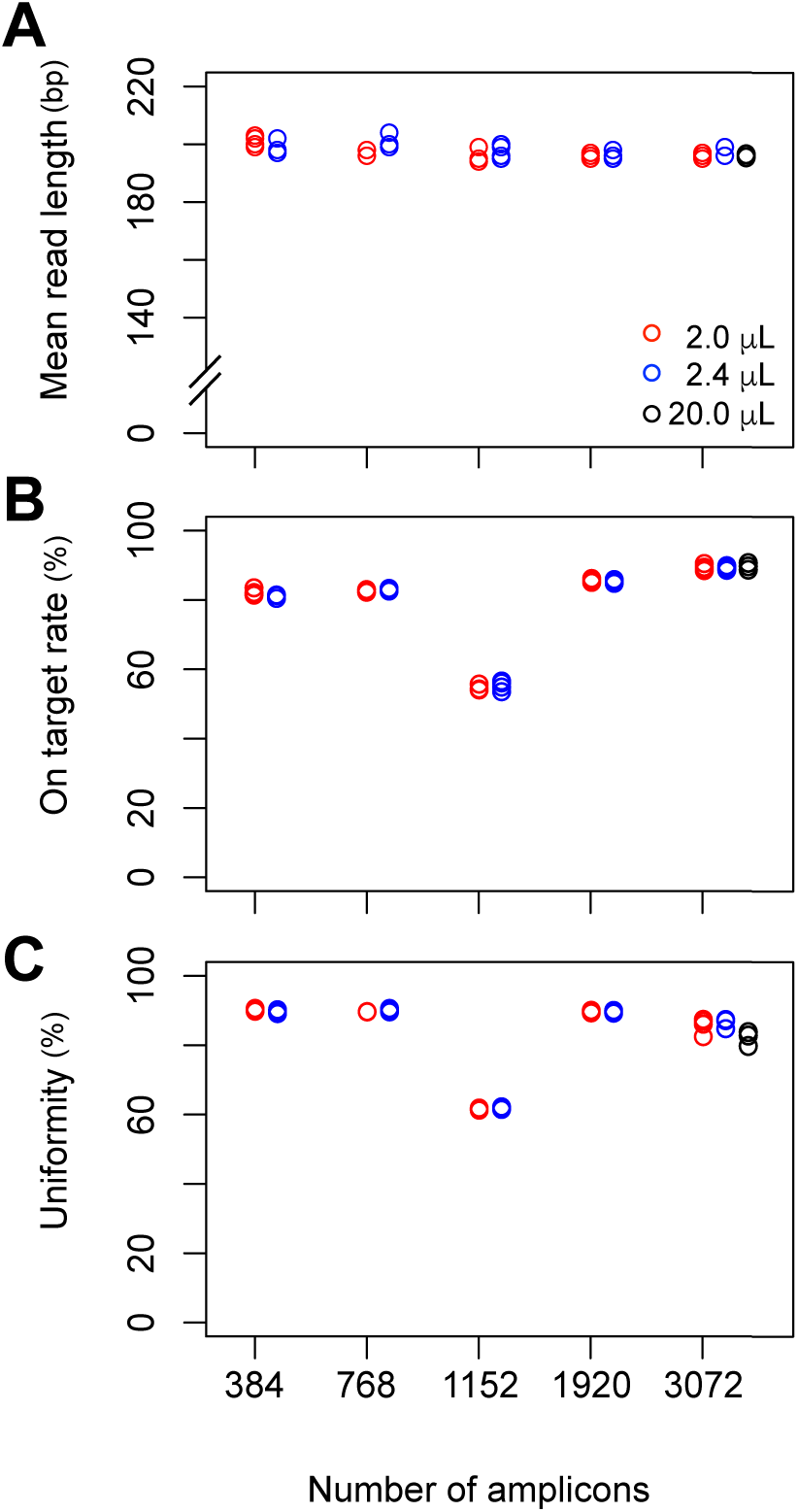
Sequencing results for four samples in 384, 768, 1152, 1920, and 3072 amplicon panels. (A) Mean read length (B) On target rate (C) Uniformity. Red, blue, and black circles indicate 2-μL, 2.4-μL, and 20-μL reaction volumes. The sequencing run and library construction for the 20-μL samples were different from those for the 2-μL and 2.4-μL samples.

### Comparison of cost and time

This miniaturized protocol reduces the cost and time for AmpliSeq library preparation. The cost of materials for each library, using the manual protocol, including consumables such as reagents, tips, plates, and seal, was 9,602 JPY (approximately 3,687,284 JPY for 384 samples), whereas the cost per sample dropped to about 1,261 JPY using this miniaturized protocol (approximately 484,325 JPY for 384 samples), thereby resulting in cost saving by about 86.8% (Table 3).

**Table 3.** Cost comparison between standard and miniaturized protocols for 384 samples

|  | Standard protocol | Miniaturized protocol |  |
| --- | --- | --- | --- |
|  | Manual |  | Mosquito HTS |
| yen [JPY] | 20 μL (100 %) | 10 μL (50 %) | 2.4 μL (12 %) |
| Reagent | 3,640,000 | 1,820,000 | 436,800 |
| Tip | 7,188 | 7,188 | 30,933 |
| AMPure | 38,592 | 19,296 | 12,864 |
| Plates/Tubes | 1,504 | 1,504 | 2,000 |
| Source Plate | N/A | N/A | 1,728 |
| Total | 3,687,284 | 1,847,988 | 484,325 |

**Table 4.** Comparison of time between manual full-volume protocol and miniaturized protocol (Mosquito HTS 2-way model) for 384 samples

|  |  | Standard protocol | Miniaturized automated protocol |  |
| --- | --- | --- | --- | --- |
| Step | min | Manual (min) | Working time (min) | Handling time (min) |
| Master mix preparation |  | 10 | - | - |
| Dispensing each reagent to source plate |  | - | 5 | 5 |
| Dispensing master mix or reagent |  | 20 | 11 | 1 |
| Multiplex PCR reaction | 74 | - | - | - |
| Dispensing FuPa solution |  | 10 | 8.2 | 1 |
| FuPa reaction | 41 | - | - | - |
| Dispensing Switch solution |  | 10 | 8.2 | 1 |
| Dispensing Barcode adaptor |  | 10 | 3.7 | 1 |
| Dispensing Ligase |  | 10 | 8.2 | 2 |
| Ligation reaction | 72 | - | - | - |
| Total | 187 | 66 min | 44.3 (67.1 %) | 12<br>(18.2 %) |

Similarly, the automated system for library preparation was time-saving. Making libraries of 384 samples by hand, at the same time, with the manual protocol, would take approximately 66 min. Although Mosquito HTS would take almost similar time (approximately 44 min) to make 384 libraries, we do not need much labor in the automated system. Since it takes time to mix reagents by pipetting, followed by individual reagent transfer (no master mix), it is difficult to shorten the library preparation time; however, the handling time can be greatly reduced due to the automated pipetting. Mosquito HTS has two models: 2-way and 5-way (2 or 5 plates can be set at once). Using the Mosquito HTS 5- way model, labor can be further minimized as some plates can be set at once and processed continuously. Alternatively, although dead volume may be generated when a master mix is prepared, dispensing time is shortened. Dead volume has little effect on the cast of library prep when dealing with sample sizes of 1000 or more.

## Discussion

In this study, we presented a miniaturization protocol for AmpliSeq library preparation. Compared to the standard protocol, our protocol is inexpensive and not labor-intensive. Empirically, stable results were obtained when the reaction volume was 2.4 μL and the number of amplicons was 1500 or less. The influence of the difference in uniformity and amplicon size of multiplex PCR was greater than the influence of the type of thermo-cycler used and reaction volume. Miniaturization seemed to make the protocol easier to be influenced by environmental factors.

This is the first application of the Mosquito HTS for the AmpliSeq ultra-multiplex PCR protocol and adds to other existing protocols for NGS library preparation [8-12]. The Mosquito HTS, used in this miniaturized and automated library preparation, enables rapid processing of a large number of samples in units of 8 (tips at 9.0-mm pitch) or 16 (4.5-mm) samples at a time, remarkably saves cost and time, and allows quick completion of large-scale genotyping.

However, there are two limitations of this Mosquito protocol. First, Mosquito HTS should be set in a cold room, whereas in this study, we used it at room temperature. Moreover, we always used freshly prepared reagents. The reagent was dispensed in advance into 384-well plate (source plate) before use, and the residual reagent was stored at −20 ºC to be used in the next experiment. As the number of times of use and time lapse increased, the quality of the library declined in the subsequent experiments (lower uniformity, data not shown). This could be prevented by using the Mosquito HTS in a cold room. Second, Mosquito HTS cannot not be used for bead clean-up. Therefore, it is necessary to use another conventional dispensing robot. As an alternative method, a two-step process was used to optimize the pooling of the samples. First, all the libraries were mixed equally and sequence-skimmed; then, the concentration from the first sequencing result was adjusted and sequenced again. Finally, the initial cost of the robot is high. However, these robots are very useful and greatly reduce the experimental cost if housed in core labs or core facilities.

In conclusion, we presented a protocol to prepare sequencing libraries in miniaturized volume using AmpliSeq library kit (Thermo Fisher Scientific, Waltham, MA). With this protocol, it is possible to prepare 384 libraries with just 12% of the standard reagent volume, at less than 13.1% of the cost, and in less than 18.2% of the time required in the standard manual protocol. This should help the advancement of not only clinical genomics, but also large-scale genotyping in the agronomic field and projects such as AgriSeq.

## Acknowledgements

The authors would like to thank Ayaka Take, Anne Hammerstein (TTP Labtech, Royston, UK), Masaya Asano, Keishi Nakayama, Masahiro Matsushita, and Naoto Momoda for their technical supports in using the Mosquito HTS and HV. This work was supported by grant from NARO, and the special scheme project on advanced research and development for next generation technology.

## References

1. Chung, Y.S., Choi, S.C., Jun, TH. et al. (2017) Genotyping-by-sequencing: a promising tool for plant genetics research and breeding. Hortic. Environ. Biotechnol. 58: 425. https://doi.org/10.1007/s13580-017-0297-8

2. Rowan B.A., Seymour D.K., Chae E., Lundberg D.S., Weigel D. (2017) Methods for Genotyping-by-Sequencing. In: White S., Cantsilieris S. (eds) Genotyping. Methods in Molecular Biology, vol 1492. Humana Press, New York, NY

3. Rasheed, A., Y. Hao, X. Xia, A. Khan, Y. Xu, R.K. Varshney, and Z. He. (2017) Crop breeding chips and genotyping platforms: progress, challenges and perspectives. Mol. Plant. doi:https://doi.org/10.1016/j.molp.2017.06.008.

4. AmpliSeq Library preparation protocol https://assets.thermofisher.com/TFS-Assets/LSG/manuals/MAN0013432_Ion_AmpliSeq_Library_Prep_on_Ion_Chef_UG.pdf

5. AgriSeq Targeted Genotyping By Sequencing https://www.thermofisher.com/jp/ja/home/life-science/agricultural-biotechnology/agrigenomics/agriseq-targeted-genotyping-sequencing.html

6. Ogiso-Tanaka E., Taguchi-Shiobara F., Hirata K., Kaga A., Hajika M., Ishimoto M. (2017). Construction of highly flexible soybean breeding panel integrating whole genome sequence and QTL information using AmpliSeq technology. Consortium of Biological Sciences (ConBio) Program: 3P–1299

7. Gaisford W, Schertler G and Edwards P (2011) mosquito LCP: Making membrane protein crystallization accessible to the research scientist. Nature Methods 8: 520

8. Mora-Castilla S, To C, Vaezeslami S, Morey R, Srinivasan S, Dumdie JN, Cook-Andersen H, Jenkins J, Laurent LC. (2016) Miniaturization Technologies for Efficient Single-Cell Library Preparation for Next-Generation Sequencing. J Lab Autom 21(4):557–67.

9. AgriSeq Targeted Genotyping By Seequencing: https://www.thermofisher.com/jp/ja/home/life-science/agricultural-biotechnology/agrigenomics/agriseq-targeted-genotyping-sequencing.html

10. Sergio Mora-Castilla, Cuong To, Soheila Vaezeslami, Robert Morey, Srimeenakshi Srinivasan, Jennifer N. Dumdie, Heidi Cook-Andersen, Joby Jenkins, Louise C. Laurent (2016) Miniaturization Technologies for Efficient Single-Cell Library Preparation for Next-Generation Sequencing. SLAS Technol. Aug; 21(4): 557–567.

11. Herrtwich, L., et. al. (2016) DNA Damage Signaling Instructs Polyploid Macrophage Fate in Granulomas. Cell, Volume 167, Issue 5, 1264–1280.e18

12. Wimmers, F., et. al. (2018) Single-cell analysis reveals that stochasticity and paracrine signaling control interferon-alpha production by plasmacytoid dendritic cells. Nature Communications 9. doi:10.1038/s41467-018-05784-.

13. Herman, J., et. al. (2018) FateID infers cell fate bias in multipotent progenitors from single-cell RNA-seq data. Nature methods. doi:10.1038/nmeth.466.

14. Sagar, et al. (2018) High-Throughput Single-Cell RNA Sequencing and Data Analysis. In: Vavouri T., Peinado M. (eds) CpG Islands. Methods in Molecular Biology, vol 1766.

15. Zanini F, Pu S, Bekerman E, Einav S, Quake S. R. (2018) Single-cell transcriptional dynamics of flavivirus infection. eLIFE 7:e32942

